# Structural basis for the recognition of the 2019-nCoV by human ACE2

**DOI:** 10.1101/2020.02.19.956946

**Authors:** Renhong Yan, Yuanyuan Zhang, Yingying Guo, Lu Xia, Qiang Zhou

## Abstract

Angiotensin-converting enzyme 2 (ACE2) has been suggested to be the cellular receptor for the new coronavirus (2019-nCoV) that is causing the coronavirus disease 2019 (COVID-19). Like other coronaviruses such as the SARS-CoV, the 2019-nCoV uses the receptor binding domain (RBD) of the surface spike glycoprotein (S protein) to engage ACE2. We most recently determined the structure of the full-length human ACE2 in complex with a neutral amino acid transporter B^0^AT1. Here we report the cryo-EM structure of the full-length human ACE2 bound to the RBD of the 2019-nCoV at an overall resolution of 2.9 Å in the presence of B^0^AT1. The local resolution at the ACE2-RBD interface is 3.5 Å, allowing analysis of the detailed interactions between the RBD and the receptor. Similar to that for the SARS-CoV, the RBD of the 2019-nCoV is recognized by the extracellular peptidase domain (PD) of ACE2 mainly through polar residues. Pairwise comparison reveals a number of variations that may determine the different affinities between ACE2 and the RBDs from these two related viruses.

## Introduction

For background introduction to ACE2 and B^0^AT1, please refer to our recent posting on *bioRxiv* that reports the 2.9 Å-resolution cryo-EM structure of the full-length ACE2 in complex with B^0^AT1, assembled as a dimer of heterodimers (1). The presence of B^0^AT1 appears to stabilize the overall conformation of the full-length human ACE2.

The surface spike glycoproteins (S proteins) of the coronaviruses are responsible for attaching to host cells through interaction with the surface receptors. S protein exists as a homotrimer, with more than 1200 amino acids in each monomer. In the S protein of the SARS-CoV, a small domain containing residues 306-575 was identified to be the receptor binding domain (RBD), in which residues 424-494 known as the receptor binding motif (RBM) directly mediate the interaction with ACE2 (2).

ACE2 has also been suggested to be the receptor for the 2019-nCoV (3, 4). The ectodomain of the 2019-nCoV S protein was reported to bind to the PD of ACE2 with a K_d_ of ~ 15 nM, measured with surface plasmon resonance (SPR) (5). Docking analysis based on our structure of the ACE2-B^0^AT1 complex suggests that RBD can access to the PD, which protrudes into the extracellular space, in the presence of B^0^AT1, indicating compatibility of all three proteins for ternary complex formation (1). Structural elucidation of the interface between S protein-RBD and ACE2-PD will not only shed light on the mechanistic understanding of viral infection, but also facilitate development of viral detection techniques and potential antiviral therapeutics. High-resolution cryo-EM structural determination of the dimer of the ACE2-B^0^AT1 heterodimers established the framework for structural resolution of the ternary complex using single-particle cryo-EM.

### Overall structure of the RBD-ACE2-B^0^AT1 complex

To reveal the recognition details between ACE2 and the 2019-nCoV, we purchased 0.2 mg recombinantly expressed and purified RBD-mFc of the 2019-nCoV (for simplicity, we will refer it as RBD for short if not specified) from Sino Biological Inc., mixed it with our purified ACE2-B^0^AT1 complex at a stoichiometric ratio of ~ 1.1 to 1, and proceeded with cryo-grid preparation and imaging. Following the same protocol for data processing as for the ACE2-B^0^AT1 complex, a 3D EM reconstruction of the ternary complex was obtained. The structural findings about the dimeric full-length ACE2 and its complex interaction with B^0^AT1, which were reported in detail in our bioRxiv posting (1), will not be repeated here. In this manuscript, we will focus on the interface between ACE2 and RBD.

In contrast to the ACE2-B^0^AT1 complex, which has two conformations, “open” and “closed”, only the closed state of ACE2 was observed in the dataset for the RBD-ACE2-B^0^AT1 ternary complex. Out of 527,017 selected particles, the overall resolution of the ternary complex was achieved at 2.9 Å. However, the resolution for the ACE2-B^0^AT1 complex is substantially higher than that for the RBDs, which are at the peripheral region of the complex (Fig. 1A). To improve the local resolutions, focused refinement was applied. Finally, the resolution of the RBDs reached 3.5 Å, supporting reliable modelling and interface analysis (Figs. 1, 2; Supplementary Figures S1,S2, Table S1).

**Figure 1.**
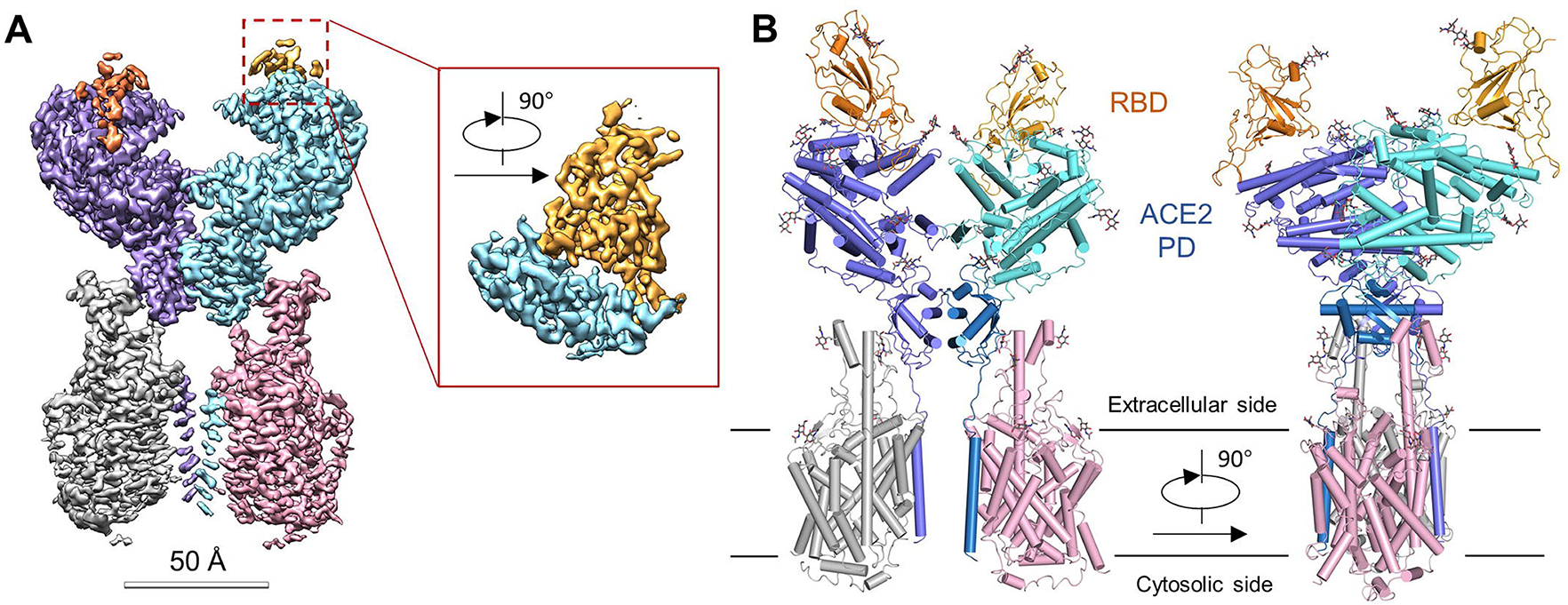
Overall structure of the RBD-ACE2-B^0^AT1 complex. **(A)** Cryo-EM map of the RBD-ACE2-B^0^AT1 complex. *Left*: Overall reconstruction of the ternary complex at 2.9 Å. *Inset*: focused refined map of RBD. (**B**) Overall structure of the RBD-ACE2-B^0^AT1 complex. The complex is colored by subunits, with the protease domain (PD) and the Collectrin-like domain (CLD) colored cyan and blue in one of the ACE2 protomers, respectively. The glycosylation moieties are shown as sticks.

**Figure 2.**
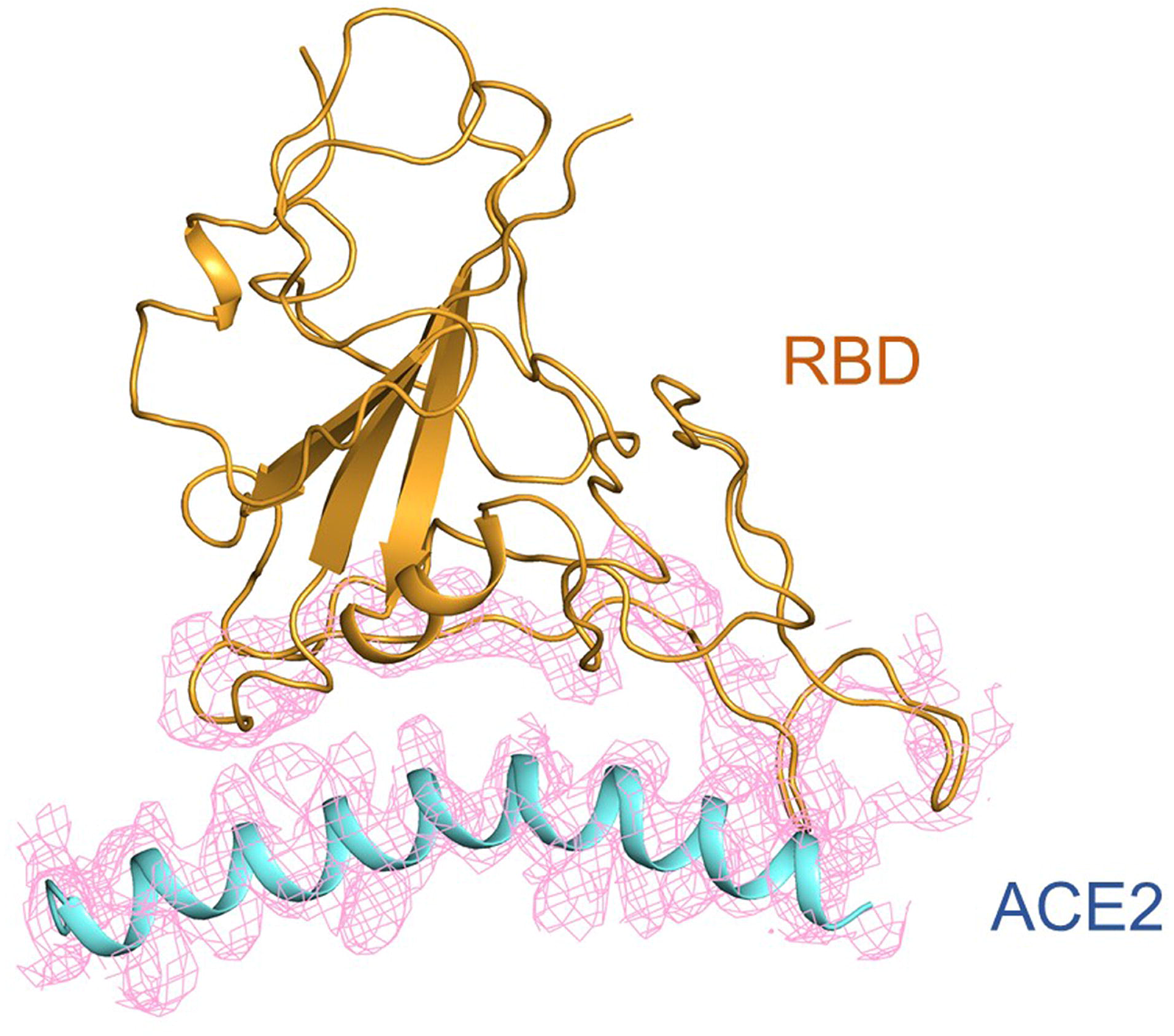
Cryo-EM density of the interface between RBD and ACE2. The density, shown as pink meshes, is contoured at 12 σ.

### Interface between RBD and ACE2

As expected, each PD accommodates one RBD (Fig. 1B). The overall interface is similar to that between the SARS-CoV and ACE2 (2, 6), mediated mainly through polar interactions (Fig. 3A). An extended loop region of RBD spans above the arch-shaped α1 helix of ACE2 like a bridge. The α2 helix and a loop that connects the β3 and β4 antiparallel strands, referred to as Loop 3-4, of the PD also make limited contributions to the coordination of RBD.

**Figure 3.**
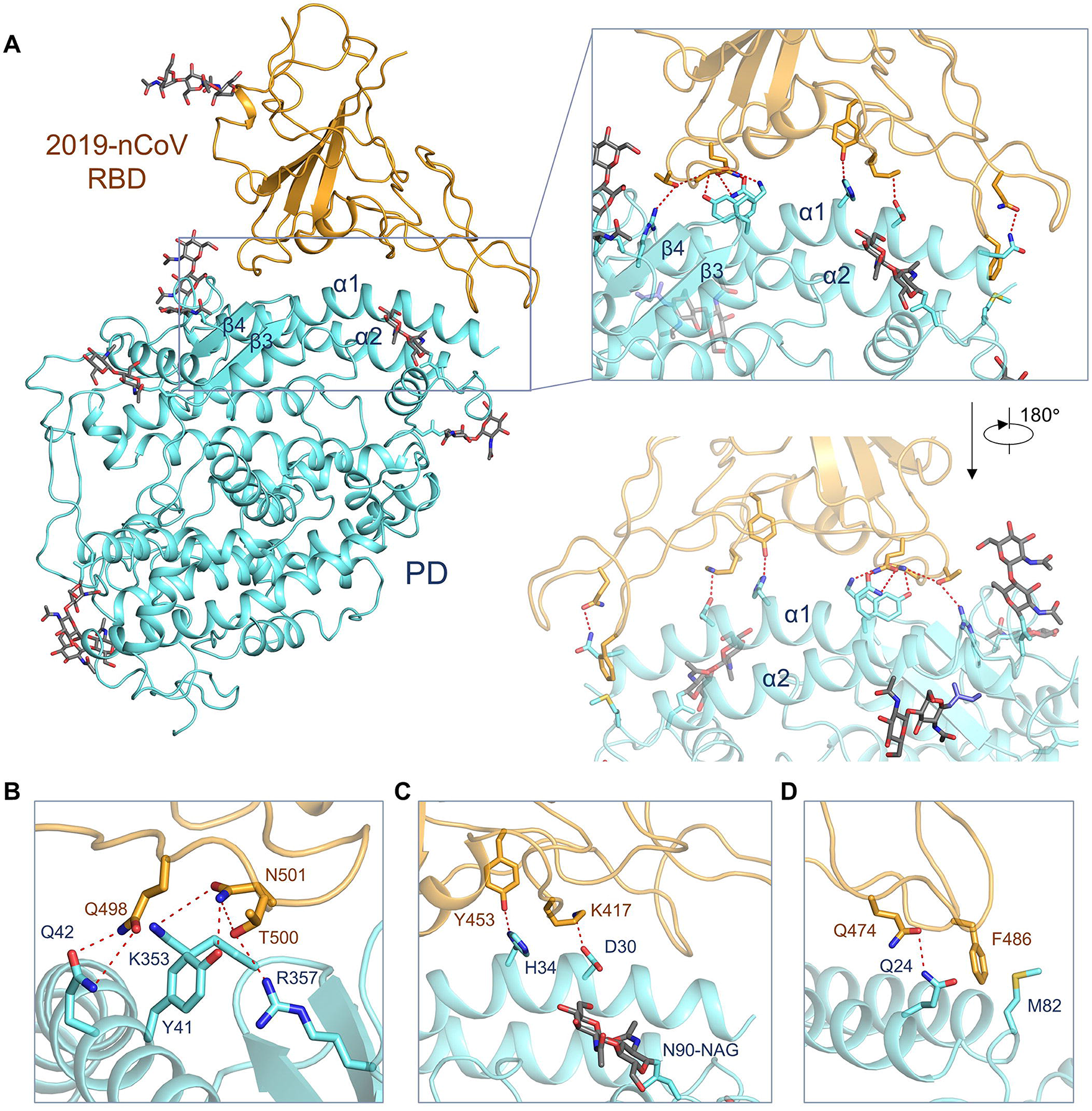
Interactions between nCoV-RBD and ACE2. **(A)** The protease domain (PD) of ACE2 mainly engages the α1 helix in the recognition of the RBD. The α2 helix and the linker between β3 and β4 also contribute to the interaction. Only one RBD-ACE2 is shown. **(B-D)** Detailed analysis of the interface between nCoV-RBD and ACE2. Polar interactions are indicated by red, dashed lines.

The contact can be divided to three clusters. The two ends of the “bridge” attach to the amino (N) and carboxyl (C) termini of the α1 helix as well as small areas on the α2 helix and Loop 3-4. The middle segment of α1 reinforces the interaction by engaging two polar residues (Fig. 3A). For illustration simplicity, we will refer to the N and C termini of the α1 helix as right and left. On the left, Gln498, Thr500, and Asn501 of the RBD form a network of hydrogen bonds (H-bonds) with Tyr41, Gln42, Lys353, and Arg357 from ACE2 (Fig. 3B). In the middle of the “bridge”, Lys417 and Tyr453 of the RBD interact with Asp30 and His34 of ACE2, respectively (Fig. 3C). On the right, Gln474 of RBD is H-bonded to Gln24 of ACE2, while Phe486 of RBD interacts with Met82 of ACE2 through van der Waals forces (Fig. 3D).

### Interface comparison between 2019-nCoV and SARS-CoV with ACE2

The structure of the 2019-nCoV RBD (nCoV-RBD) is similar to the RBD of SARS-CoV (SARS-RBD) with a root mean squared deviation of 0.68 Å over 139 pairs of Cα atoms (Fig. 4A) (2). Despite the overall similarity, a number of sequence variations and conformational deviations are found on their respective interface with ACE2 (Fig. 4, Supplementary Figure S3). On the left end of the “bridge”, Arg426 →Asn439, Tyr484 → Gln498, and Thr487 → Asn501 are observed from SARS-RBD to nCoV-RBD (Fig. 4B). More variations are observed in the middle of the bridge. The most prominent alteration is the substitution of Val404 in the SARS-RBD with Lys417 in the nCoV-RBD. In addition, from SARS-RBD to nCoV-RBD, the following alternation of interface residues, Tyr442 → Leu455, Leu443 → Phe456, Phe460 → Tyr473, and Asn479 → Gln493, may also change the affinity with ACE2 (Fig. 4C). On the right end, the corresponding locus for Leu472 in the SARS-RBD is occupied by Phe486 in the nCoV-RBD (Fig. 4D).

**Figure 4.**
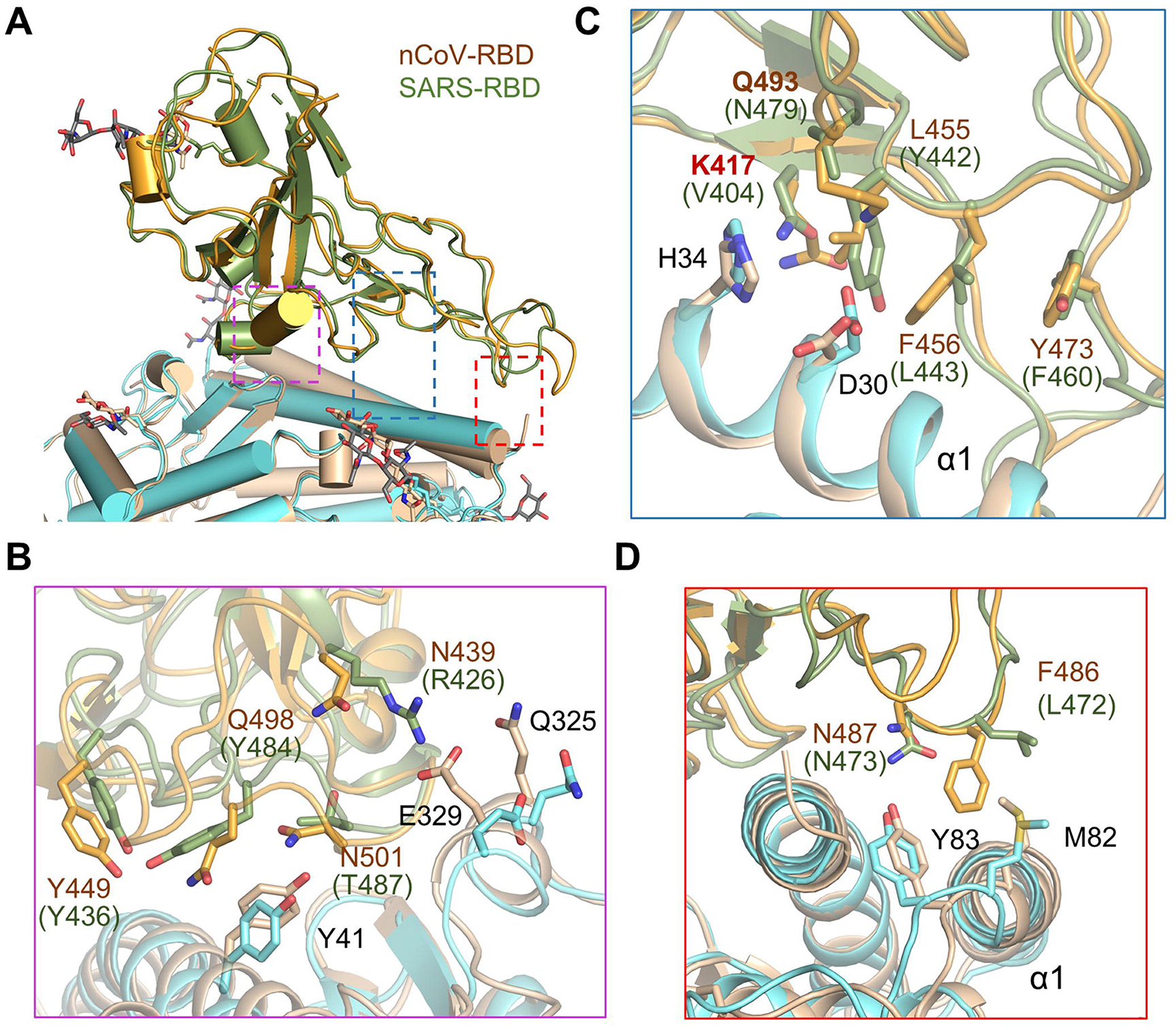
Interface comparison between nCoV-RBD and SARS-RBD with ACE2. **(A)** Structural alignment for the nCoV-RBD and SARS-RBD. The complex structure of ACE2 and SARS-RBD (PDB code: 2AJF) is superimposed to our cryo-EM structure. The boxed regions are illustrated in details in panels ***B-D***. nCoV-RBD and the PD in our cryo-EM structure are coloured orange and cyan, respectively; SARS-RBD and its complexed PD are coloured green and gold, respectively. **(B-D)** Variation of the interface residues between nCoV-RBD (labeled brown) and SARS-RBD (labeled green). In this three panels, the two structures are superimposed relative to RBD.

## Discussion

On the basis of our cryo-EM structural analysis of the ACE2-B^0^AT1 complex, we hereby report the high-resolution cryo-EM structure of the ternary complex of RBD from the 2019-nCoV associated with full-length human ACE2 in the presence of B^0^AT1.

The 2019-nCoV has killed more people in the past two months than SARS-CoV because of its high infectivity, whose underlying mechanism remains unclear. A furin cleavage site unique to the S protein of the 2019-nCoV may contribute to its greater infectivity than SARS-CoV (7, 8). The recently reported higher affinity between the nCoV-RBD and ACE2 may represent an additional factor (5).

Structures of ACE2 in complex with nCoV-RBD and SARS-RBD establish the molecular basis to dissect their different affinities. In this study, we carefully analyzed the interface in both RBD-ACE2 complexes. Whereas some of the variations may strengthen the interactions between nCoV-RBD and ACE2, others may reduce the affinity compared to that between SARS-RBD and ACE2. For instance, the change from Val404 to Lys317 may result in tighter association because of the salt bridge formation between Lys317 and Asp30 of ACE2 (Figs. 3C, 4C). Change of Leu472 to Phe486 may also make stronger van der Waals contact with Met82 (Fig. 4D). However, replacement of Arg426 to Asn439 appears to weaken the interaction by losing one important salt bridge with Asp329 on ACE2 (Fig. 4B).

Our structure provides the molecular basis for computational and mutational analysis for the understanding of the affinity difference. It should be noted that additional approaches, such as isothermal titration calorimetry (ITC) and microscal thermophoresis (MST), should be applied to validate SPR-measured affinities.

Structural elucidation nCoV-RBD to ACE2 also set up the framework for the development of novel viral detection methods and potential therapeutics against the 2019-nCoV. Structure-based rational design of binders with enhanced affinities to either ACE2 or the S protein of the coronaviruses may facilitate development of decoy ligands or neutralizing antibodies that block viral infection.

## Acknowledgments

We thank the Cryo-EM Facility and Supercomputer Center of Westlake University for providing cryo-EM and computation support, respectively. This work was funded by the National Natural Science Foundation of China (projects 31971123, 81920108015, 31930059) and the Key R&D Program of Zhejiang Province (2020C04001).

## Author contributions

Q.Z. and R.Y. conceived the project. Q.Z and R.Y. designed the experiments. All authors did the experiments. Q.Z. R.Y., and Y.Z. contributed to data analysis. Q.Z. and R.Y. wrote the manuscript.

