## Supplementary for "Structural basis for the recognition of the 2019-nCoV by human ACE2"

### Methods

#### Cryo-EM sample preparation and data acquisition

The purified ACE2-$B^0AT1$ complex was mixed with the RBD (residues 319-541) of the S protein of the 2019-nCoV, with a C-terminal mouse Fc tag (Sino Biological Inc.) at a molar ratio of about 1:1.1 in the presence of 10 mM leucine. After 2 hour-incubation, aliquots (3.3 μl) of the mixture were placed on glow-discharged holey carbon grids (Quantifoil Au R1.2/1.3), which were blotted for 3.0 s or 3.5 s and flash-frozen in liquid ethane cooled by liquid nitrogen with Vitrobot (Mark IV, Thermo Fisher Scientific). The cryo grids were transferred to a Titan Krios operating at 300 kV equipped with Gatan K3 Summit detector and GIF Quantum energy filter. Movie stacks were automatically collected using AutoEMation (*1*), with a slit width of 20 eV on the energy filter and a defocus range from -1.2 μm to -2.2 μm in super-resolution mode at a nominal magnification of 81,000 ×. Each stack was exposed for 2.56 s with an exposure time of 0.08 s per frame, resulting in a total of 32 frames per stack. The total dose rate was approximately 50 $e^-/Å^2$ for each stack. The stacks were motion corrected with MotionCor2 (*2*) and binned 2-fold, resulting in a pixel size of 1.087 Å/pixel. Meanwhile, dose weighting was performed (*3*). The defocus values were estimated with Gctf (*4*).

#### Data processing

Particles were automatically picked using Relion 3.0.6 (*5-8*) from manually selected micrographs. After 2D classification with Relion, good particles were selected and subject to three cycles of heterogeneous refinement with C1 symmetry using cryoSPARC (*9*). The good particles were selected and subject to homogeneous refinement with C2 symmetry, resulting in a 3D reconstruction at 2.9 Å. To further improve the map quality of RBD, the particles were C2-symmetry expanded and re-extracted at the location of the interface between ACE2 and RBD. The re-extracted dataset was focused 3D classified with Relion using the alignment parameters output from cryoSPARC. Then the good particles were selected and subject to focused refinement with Relion, resulting in a 3D reconstruction with better quality for RBD. The resolution was estimated with the gold-standard Fourier shell correlation 0.143 criterion (*10*) with high-resolution noise substitution (*11*). Please refer to Supplementary Figures S1-S2 and Table S1 for details of data collection and processing.

**Model building and structure refinement**

The model building for the part of the ACE2- $B^0$AT1 complex was accomplished with Phenix (*12*) and Coot (*13*) based on the cryoSPARC map using the model of the ACE2-$B^0$AT1 complex described previously as initial template. The atomic model of the SARS-RBD (PDB ID: 2AJF) was sequence-substituted to the nCoV-RBD in chainsaw and fitted into the focused refined map of RBD using MDFF (molecular dynamics flexible fitting) (*14*). Each residue was manually checked with Coot with the chemical properties taken into consideration during model building. Statistics associated with data collection, 3D reconstruction and model building is summarized in Supplementary Table S1.

**Supplementary Table S1 | Data collection, 3D reconstruction and model statistic**

| | | |
|---|---|---|
| **Data collection** | | |
| EM equipment | Titan Krios (Thermo Fisher Scientific) | |
| Voltage (kV) | 300 | |
| Detector | Gatan K3 Summit | |
| Energy filter | Gatan GIF Quantum, 20 eV slit | |
| Pixel size (Å) | 1.087 | |
| Electron dose (e-/Å2) | 50 | |
| Defocus range (μm) | -1.2 ~ -2.2 | |
| Number of collected micrographs | 8,055 | |
| Number of selected micrographs | 7,532 | |
| Conformation | closed, RBD bound | interface of RDB and ACE2 |
| **3D Reconstruction** | | |
| Software | cryoSPARC | Relion |
| Number of used particles | 527,017 | 301,565 |
| Resolution (Å) | 2.9 | 3.5 |
| Symmetry | C2 | C1 |
| Map sharpening B factor (Å2) | -118 | -150 |
| **Refinement** | | |
| Software | Phenix | |
| Cell dimensions | | |
| a=b=c (Å) | 313.056 | |
| α=β=γ (˚) | 90 | |
| Model composition | | |
| Protein residues | 3,078 | |
| Side chains assigned | 3,078 | |
| Sugar | 44 | |
| ligand | 6 | |
| substrate, Leu | 2 | |
| Zn | 2 | |
| Water | 8 | |
| R.m.s deviations | | |
| Bonds length (Å) | 0.008 | |
| Bonds Angle (˚) | 1.187 | |
| Ramachandran plot statistics (%) | | |
| Preferred | 91.01 | |
| Allowed | 8.53 | |
| Outlier | 0.46 | |

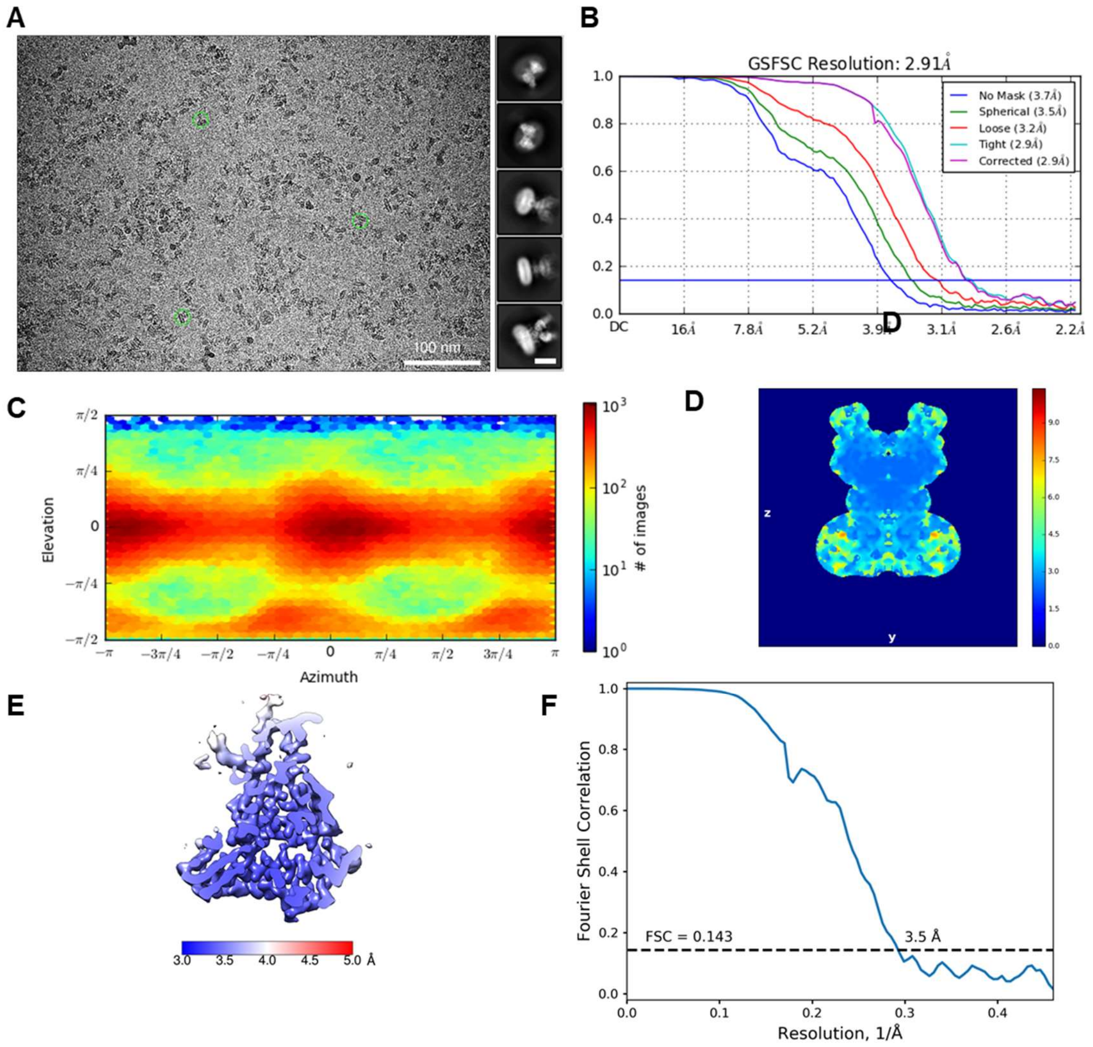


**Supplementary Figure S1**

Cryo-EM analysis of the RBD-ACE2-$B^0$AT1 ternary complex.

**(A)** Representative electron micrograph and 2D class averages of cryo-EM particle images. The scale bar in 2D class averages represents 10 nm. **(B)** Gold standard FSC curve of the cryoSPARC 3D reconstruction of the ternary complex. **(C)** Euler angle distribution of the ternary complex in the final cryoSPARC 3D reconstruction. **(D)** Local resolution map for the 3D reconstruction of the complex. **(E)** Local resolution map for the 3D reconstruction of the interface between RBD and ACE2. **(F)** Gold standard FSC curve of the interface between RBD and ACE2.

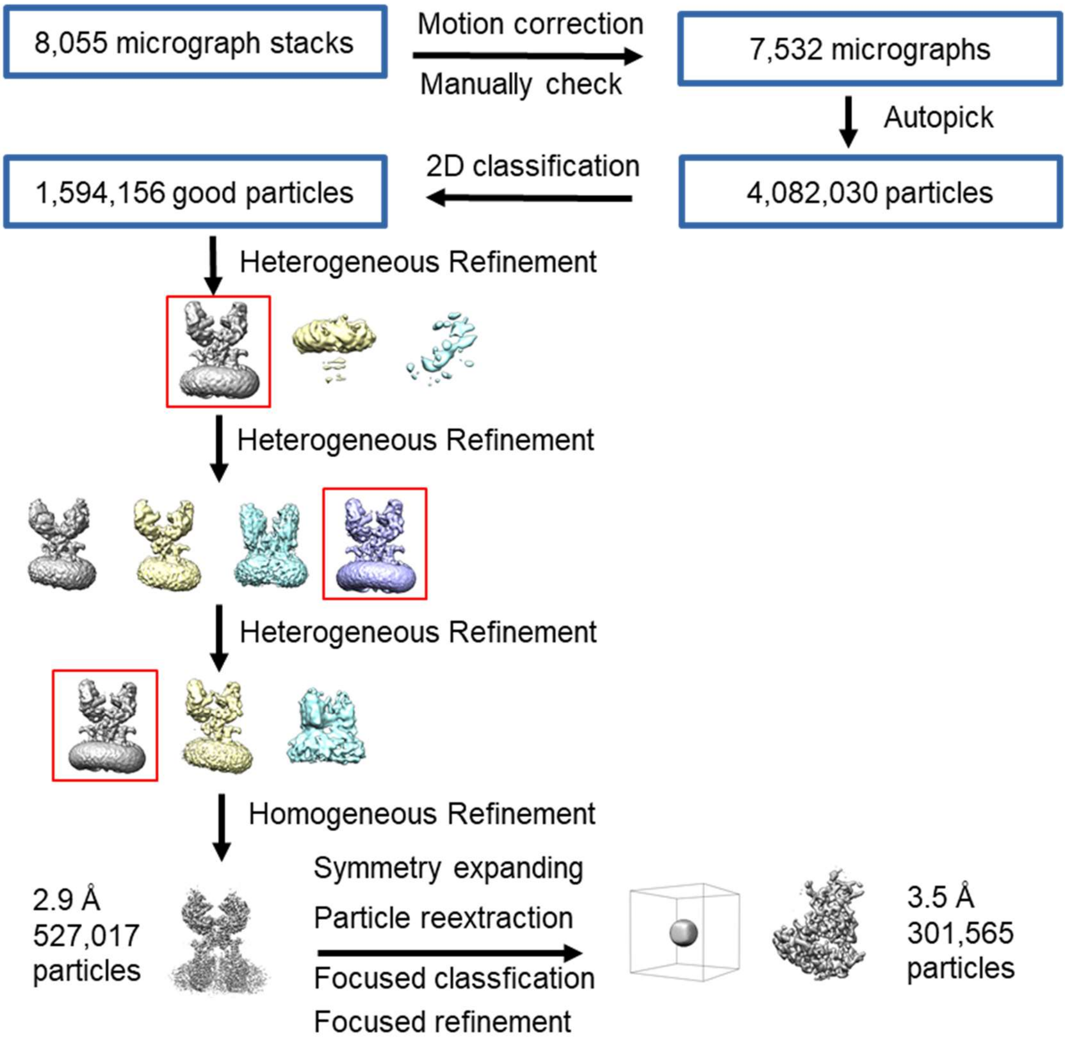


**Supplementary Figure S2**

Flowchart for cryo-EM data processing.

Please see the "Data Processing" section in Methods for details.

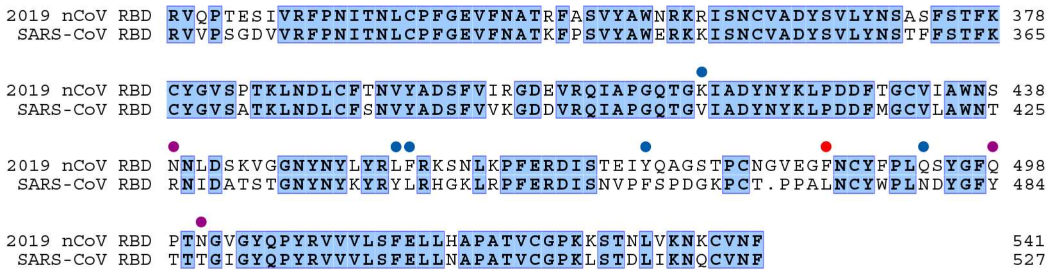


**Supplementary Figure S3**

Sequence alignment for the RBD of the S proteins from 2019-nCoV and SARS-CoV.

The two sequences were aligned using ClustalX. Invariant amino acids are shaded blue. The altered interface residues between nCoV-RBD and SARS-CoV RBD are indicated by solid circles and color-coded using the same scheme as for the boxes in Figure 4. The Uniprot IDs: 2019-nCoV S protein (P0DTC2) and SARS-CoV S protein (P59594).
